# D2Sim: A Computational Simulator for Nanopore Sequencing based DNA Data Storage

**DOI:** 10.1101/2024.03.17.585393

**Authors:** Subhasiny Sankar, Wang Yixin, Md. Noor-A-Rahim, Erry Gunawan, Yong Liang Guan, Chueh Loo Poh

## Abstract

DNA data storage has gained significant attention because of its high storage density and durability. However, errors during the storage and reading processes compromise data integrity, prompting research into error correction strategies. Researchers have explored physical (data copies) and logical redundancy (added redundancy in error-correcting codes) to mitigate errors. Evaluating these designs and reconstruction methods typically involves time-consuming and costly trials. To streamline this process, we designed a computational channel simulator, namely D2Sim, for Nanopore sequencing-based DNA data storage. This simulator mimics real experiments, generating data with certain redundancy and errors at the receiver. Integrated with DeepSimulator, D2Sim outputs signals closely resembling actual signals of Nanopore-based DNA storage. Comparative analysis reveals that the proposed simulator yields 16.7% to 88.7% lower sample difference deviations than signals from DeepSimulator alone. This cost-effective and time-efficient tool facilitates the assessment of physical and logical redundancy for data reconstruction in DNA data storage without the need for real-time experiments.

## I. Introduction

**I**N DNA data storage technology, the data is encoded, synthesized/written in the form of DNA molecules, amplified by polymerase chain reaction (PCR) for sequencing/reading data from DNA molecules using commercial sequencers such as Illumina or Nanopore platform, and decoded back to the original data (Fig. 1). After writing, storing, and retrieving, at the receiver, the original encoded sequences might get lost (termed as sequence dropout) and corrupted by errors due to insertion, deletion, and substitution of molecules. Sequencing with Illumina/Nanopore is the major contributor to these errors and losses [1]. There are two ways to recover the data, that is, using physical and/or logical redundancy [2]. Physical redundancy, also termed sequencing coverage, refers to DNA copies extracted at the sequencing stage, and logical redundancy is the redundancy added while encoding with error control codes, which were later used to correct the errors at the decoder.

**Fig. 1.**
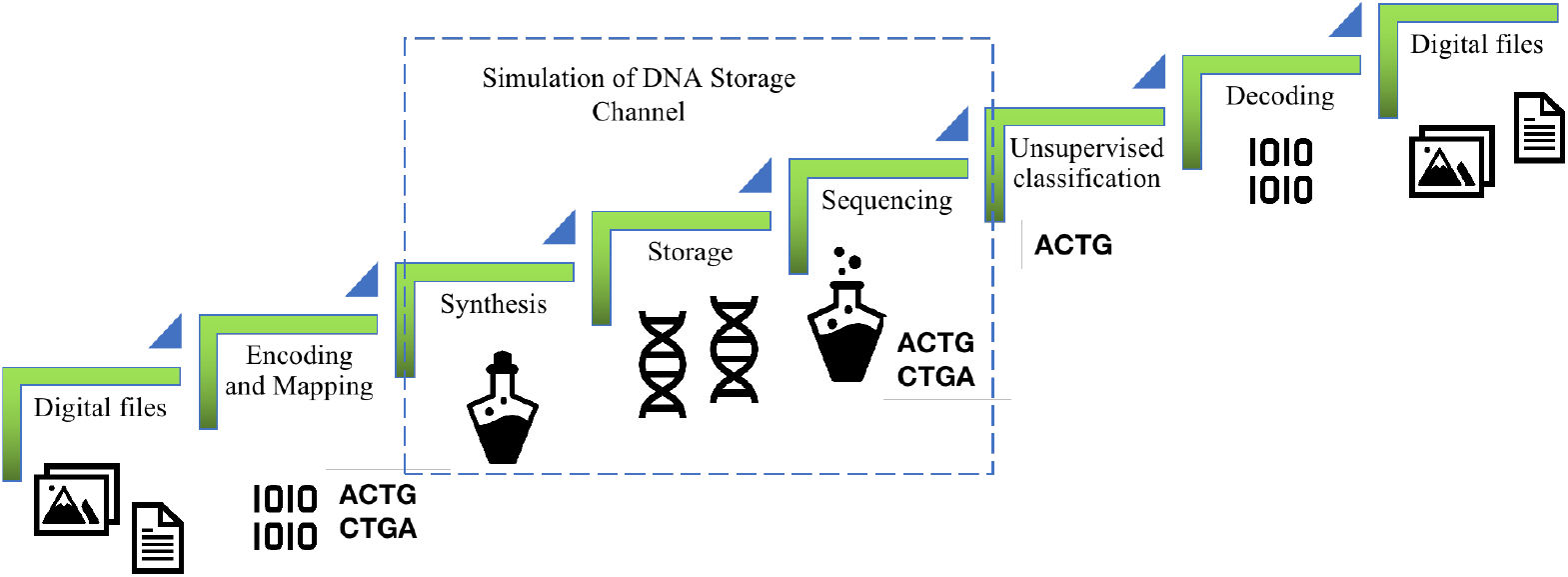
DNA storage architecture: Data is encoded, mapped to DNA, passed to DNA storage channel (represented in dotted rectangle), clustered, and decoded back to real data. Simulating DNA storage channel gives insights to decide the rest of the data flow.

Several research groups have predicted the sequencing coverage required through real-time experiments [3]–[10]. The Illumina sequencing technique, known for its lower cost and error rate of just 1%, was preferred predominantly before the advent of Nanopore sequencing. Researchers predicted sequencing redundancy levels based on factors such as the size of encoded data, the coding methods used [11], clustering techniques to group all sequences originating from the same encoded sequences, data reconstruction method to determine encoded sequence before decoding [12] [13], and the logical redundancy incorporated [3]–[For instance, Erlich et al. introduced a DNA data storage method using fountain codes, which showed that deep copy extraction and sampling sub-sequences result in a skewed negative binomial distribution. Their experiments indicated that with a size parameter of r = 7 and a coverage size of 7, a low drop-out rate of 0.5% can be achieved [3].

Recently, Nanopore sequencing has gained significant attention due to its portability and ability to read long DNA strands, making it particularly suitable for DNA data storage applications. In this process, DNA strands pass through a nanopore, generating an electrical signal for every k successive nucleotides, known as k-mers. A basecaller then translates these electrical signals into nucleotide/basecalled sequences. A study conducted during the early access program of Nanopore revealed that the reads exhibited a total error rate of 10.5% [14]. This highlights the challenges associated with the technology, particularly its high error rates [15]. Fortunately, setting up higher sequencing redundancy can compensate for these high error rates. Existing DNA data storage systems that utilize Nanopore sequencing require a sequencing coverage ranging from 75 to 566 for complete data recovery, as reported in one study [8], and from 74 to 147 in another study [5]. This variability in coverage highlights the challenges associated with accurately retrieving data from DNA storage systems. There is also a study with a fragment length-increasing method that utilizes a DNA assembly strategy to construct large DNA fragments from short DNA, and it has successfully reduced the coverage requirement to 22 [9]. The authors in [10] performed inner decoding on intermediate transition probabilities obtained from the Flappie base caller. With an average of 4 copies per sequence, they successfully decoded the data back with outer decoding. These advancement underscores the continuous efforts to enhance the efficiency and effectiveness of DNA data storage solutions by minimizing sequencing redundancy and enabling accurate data recovery, even in the presence of high error rates.

In summary, independent trial experiments have been conducted to explore the minimum redundancy required for data recovery in DNA data storage with different system settings, incurring significant costs in time and expense [5], [8], [16]. The statistical errors leading to the loss of sequences make data redundancy a necessity for perfect data retrieval [17]. The DNA that has fed into the sequencing experiment should have sufficient data variability based on a certain distribution. These explorations and validations could be replaced with an experimental simulator, saving time, cost, and reducing the number of runs. The statistical insights of distribution plays an important role in designing optimal error correction coding. To the best of our knowledge, three groups [18] [19] [20] introduced a statistical model for DNA data storage using Illumina sequencing. The first two works [18] [19] included the modelling of the distribution for error rate and coverage bias. And another work [20] uses a Generative Adversarial Network (GAN) model to train the experimental unique input sequences and many output sequences for matching their distribution of errors. The actual error statistics are reported by the model.

We developed a specialized model tailored for nanopore sequencing-based DNA data storage. This model serves as a benchmark for evaluating novel encoding, decoding, and data reconstruction techniques, addressing the unique challenges posed by nanopore sequencing technology. While there are biological simulators available for simulating Nanopore sequencing [21]–[23], they lack the option of tuning sequencing coverage, and they don’t include synthesis and PCR phase, which is highly required in the DNA storage domain. Sequencing coverage plays a significant role in assessing redundancy and factoring error control coding design. Therefore, designing a computational simulator model that allows for the tuning of sequencing coverage is sought after.

While several simulators exist to mimic the nanopore sequencing process, only two simulators have been specifically designed to model the entire DNA storage channel based on nanopore sequencing. The study, by [24], modeled substitution and insertion errors as uniformly distributed, while deletion errors were modeled as tail-favored during the synthesis phase. To account for the shortening of DNA lifespan due to environmental factors like temperature and humidity, the storage phase was modeled using the Arrhenius equation. PCR amplification was simulated by fetching a uniform random sample after each cycle, with bias toward specific samples diminishing over successive cycles. In the sequencing phase, errors were again modeled as uniformly distributed, similar to the synthesis phase. The study compared experimental errors with simulated errors by mapping encoded sequences to their sequenced or simulated outputs. Additionally, the reconstruction efficiency of experimental decoded datasets was evaluated.

Another study, by [25], introduced a statistical DNA storage channel model based on a Markov chain and proposed a non-binary LDPC codes consensus algorithm for data reconstruction. However, they identified a limitation in reproducing experimental channel behavior using DeepSimulator, as it generated the same type of error across all sequences. The designed Markov chain model incorporated the statistical dependency between input (X) and output (Y) with memory similar to DeepSimulator, and they reported that the error distribution closely matched the experimental channel. However, these two studies [24], [25] quantifies error rate but did not consider coverage biases.

Loss of sequences due to uneven coverage distribution is a significant challenge in DNA data storage systems, despite their high storage density. This issue is more critical than errors caused by mutations, as it directly impacts the reliability of data reconstruction. Sequencing redundancy, or coverage, plays a vital role in addressing this challenge, as it aids in designing robust coding and reconstruction techniques. To tackle this, we developed a new simulator, D2Sim, which introduces the ability to specify sequencing coverage and simulate sequences accordingly. D2Sim incorporates a synthesizing/PCR error module and a distribution generation module that simulates sequencing bias distributions based on actual Nanopore DNA storage data. This module employs a data-driven design of redundancy and a non-parametric subsampling method to generate realistic distributions. By integrating these features with DeepSimulator [26], D2Sim overcomes the limitation of generating identical errors across all sequences, as noted in prior studies [25], and provides a more accurate and flexible simulation framework for DNA data storage applications.

Overall, D2Sim provides a comprehensive tool for modeling the DNA storage channel, encompassing the writing, storing, and reading processes. This computational simulator could be used for evaluating the performance of DNA data storage systems by exploring optimal coding and system designs that balance logical and physical redundancy. Additionally, we validated D2Sim by comparing simulated and real nanopore signals and reads at both the electrical signal level and the basecalled sequence level.

## II. Materials and Methods for Designing Proposed Simulator

### A. Difference of simulator requirement in biological and DNA data storage domain

In genomic sequencing, the long genome/DNA sequence is split into several overlapping and non-overlapping sequences. The length of these genome segments varies and is determined by a factor called coverage (number of reads) and the distribution of read lengths. To study the distribution of read lengths, as well as error distribution and error patterns, a biological simulator that mimics the sequencing process is designed. However, in the case of DNA data storage technology, coverage is specified as the ratio of the number of sequence readouts to the number of encoded original sequences [7], which differs from the coverage concept used in biological simulator tools [21].

Several simulators are designed to replicate the processes of Nanopore sequencers within the biological domain, including Silico, Nanosim, and DeepSimulator. Both Silico [22] and Nanosim [23] utilize predefined statistical error models that are based on previously observed empirical datasets. However, these models may not be optimal, as their performance can vary significantly depending on the basecaller used to convert the raw signal into a nucleotide sequence after sequencing. This variability highlights the importance of selecting a simulator that aligns well with the specific characteristics of the sequencing technology employed.

DeepSimulator models the sequencing process by generating signals that mimic those produced by Nanopore sequencers and subsequently basecalling these signals using standard basecallers to produce nucleotide sequences. Due to its ability to approximate the sequencing process and provide outputs in multiple forms—both raw signals and nucleotide sequences—DeepSimulator is widely regarded as a valuable tool for simulating DNA data storage workflows. However, DeepSimulator focuses exclusively on simulating the sequencing process and does not account for other critical steps in DNA data storage, such as DNA synthesis or PCR amplification, where the number of sequences is amplified exponentially. Additionally, it lacks features for customizing data distribution based on sequencing coverage, which may limit its flexibility for DNA storage applications.

Simulators generate noisy reads using model error profile from experimental data. The errors are introduced by modifying bases or introducing errors in the basecaller stage while translating signals into nucleotides. These error models are from complete genomes and the length is longer than synthetic DNA, which is limited to 300nt.

### B. DeepSimulator

DeepSimulator [21] [26] is a simulation tool designed to replicate the sequencing process of Nanopore sequencers, consisting of three main components: a sequence generator, a signal generator, and a basecaller. In the context of DNA data storage, the preprocessing stage of the sequence generator has been modified to sample full-length sequences with specified coverage, rather than short sequences based on read length distribution performed in biological data [24]. The signal generator simulates the electrical signals produced during sequencing and includes two types of pore models. The context-dependent pore model operates using a bidirectional Long Short-Term Memory (bi-LSTM) network, which is trained on biological datasets to effectively capture the sequential dependencies of nucleotide signals. In contrast, the context-independent pore model is based on the official Nanopore k-mer model, making it particularly suitable for DNA data storage applications. This model allows for customization and variation that diverges from traditional biological data, accommodating the unique characteristics of synthetic DNA. In the context-independent pore model, sequences are transformed into event sequences, where each 6-mer is mapped to a corresponding current signal. This mapping is achieved using a Gaussian distribution characterized by a specified mean and standard deviation, ensuring that the simulated signals accurately reflect the expected electrical responses during sequencing.

Additionally, the signal generator features a signal repeating component that ensures the simulated signals accurately reflect the repetitive nature of Nanopore sequencing signals. This component modifies the current signal or event sequence into a ground truth signal by matching the scale of the simulated signal to resemble Nanopore reads. It does so by repeating the signal at each position based on a signal repeat time distribution (computed from Deep Canonical Time Warping with datasets), which also introduces noise into the simulated signal. And then, the signal is processed through a low-pass filter to remove high-frequency components embedded in square waves, resulting in a signal that resembles Nanopore signals. Gaussian noise is added at each position of the simulated signal using parameters for raw signal noise for event-level noise, allowing for variability in the signal for the same 6-mer at different positions. These features collectively enhance the realism of the simulated signals, improving their utility for downstream basecalling and analysis. The basecaller module translates the simulated signals into nucleotide sequences using neural network-based basecallers such as Albacore, Guppy, and Flappie.

While DeepSimulator effectively simulates the sequencing process, it has limitations when applied to DNA data storage, as it does not simulate other critical steps, such as DNA synthesis or PCR amplification, where sequences are exponentially amplified, and it lacks the ability to customize data distribution based on sequencing coverage. To address these limitations, we developed a new simulator, D2Sim, which integrates additional features into DeepSimulator to better suit DNA data storage scenarios. A synthesizing/PCR error module was added to simulate the overall error introduced during these processes, along with a distribution generation module that simulates sequencing bias distributions based on actual Nanopore DNA storage data as highlighted with dotted box in (Fig. 2). By integrating these modules with DeepSimulator, D2Sim provides a more comprehensive simulation framework tailored to the unique requirements of DNA data storage applications. We have also integrated the Flappie basecaller [27] for validation since the real Nanopore signals and basecalled sequences for comparison are from [10] and they used the Flappie basecaller. Moreover, we have maintained a 10.5% error rate, which is similar to real Nanopore signals. The primary objective of this section is to explain the functionality of DeepSimulator, particularly its pore model and signal repeating component, and to highlight the modifications made to create D2Sim, which addresses the limitations of existing simulators, enabling more accurate and flexible simulations for DNA data storage applications.

**Fig. 2.**
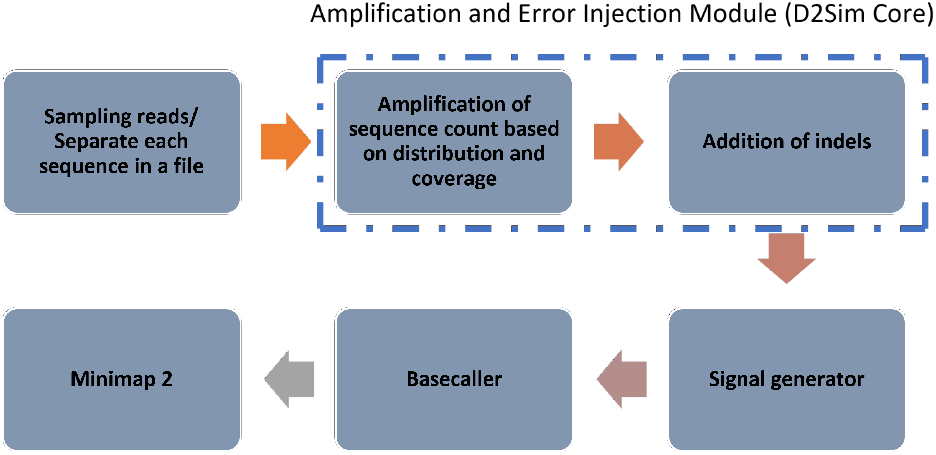
Overview of the D2Sim signal simulation pipeline. Input sequences are first sampled and separated into individual files. The amplification and error injection module (dotted rectangle) simulates amplification based on the modeled coverage distribution and inserts indel errors to mimic synthesis/PCR effects. The modified sequences are then passed to a signal generator, followed by nanopore basecalling. Finally, the resulting reads are aligned back to the original sequences using Minimap2 to assess error profiles and recovery performance.

### C. Temporal Alignment Validation

To assess the temporal alignment between the simulated signals and real Nanopore experimental signals, Dynamic Time Warping (DTW) is applied to temporally align signals, allowing for nonlinear, elastic shifts in time. This step ensures a more robust alignment compared to direct linear matching, especially when signals differ slightly in length or temporal structure.

Let *X* = [*x*_1_, *x*_2_, …, *x*_*N*_] and *Y* = [*y*_1_, *y*_2_, …, *y*_*M*_] represent two time series of length *N* and *M*, respectively. DTW computes the optimal warping path that minimizes the cumulative distance between them. The cost matrix *D*(*i, j*) is defined recursively as:

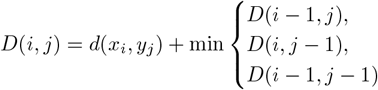

with the local distance function defined as:

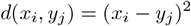

Boundary conditions:

- *D*(0, 0) = 0
- *D*(*i*, 0) = *D*(0, *j*) = *∞* for *i, j >* 0

For reproducibility and clarity, a full worked example of the DTW application with equations and signal illustrations is provided in the Supplementary Material.

After alignment using DTW, the resulting time-warped signals, *x*_DTW_ and *y*_DTW_, were used to compute the sample difference deviation (SDD), defined as the number of time samples required to best align peaks or structural features between the two sequences. In other words, it is defined as the lag (in samples) at which the cross-correlation between the aligned signals reaches its maximum absolute value:

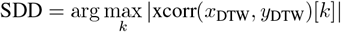

A lower sample difference deviation indicates a better temporal match between the simulated output and the real Nanopore signal, highlighting the simulator’s ability to preserve temporal fidelity.

## II. Proposed Variant of Simulator

We propose a DNA data storage simulator, D2Sim, designed to replicate the processes of synthesizing, amplifying, and sequencing DNA data (Fig. 2). The simulator generates multiple copies of each encoded DNA sequence, reflecting the distribution of sequence copies that would naturally result from these processes. The copy count refers to the number of times a specific sequence is represented after amplification and sequencing, which varies due to the stochastic nature of these processes. To estimate this copy count distribution, we employ a non-parametric subsampling method [28], [29], which allows us to model the variability in the number of copies for each sequence based on real-world data distributions.

In addition to modeling copy counts, the simulator embeds errors such as insertions, deletions (indels), and substitutions into the sequences at specified rates, simulating the types of errors introduced during the DNA synthesis and PCR process. These errors are randomly applied to the generated copies according to the error rate, ensuring a realistic representation of the challenges encountered in DNA data storage. The design of D2Sim follows five key steps.

### A. Detection of distribution

We estimated the copy distribution of the sequenced data from each experiment conducted in [10] by mapping the sequenced reads with reference IDs. Using data obtained from 11 experiments, we conducted an analysis to determine the probabilistic model associated with the channel. We evaluated the derived distribution using criteria including Sum of Squared Errors (SSE), Akaike Information Criterion (AIC), and Bayesian Information Criterion (BIC) (Fig. 3a). Mostly, the distributions observed in the 10 experiments are normal, beta, and log-gamma distributions (Supplementary S1 Table 1). For modeling these distributions after evaluating the best probabilistic model, we compared two sampling methodologies, such as parametric and non-parametric and so the better method is determined.

**Fig. 3.**
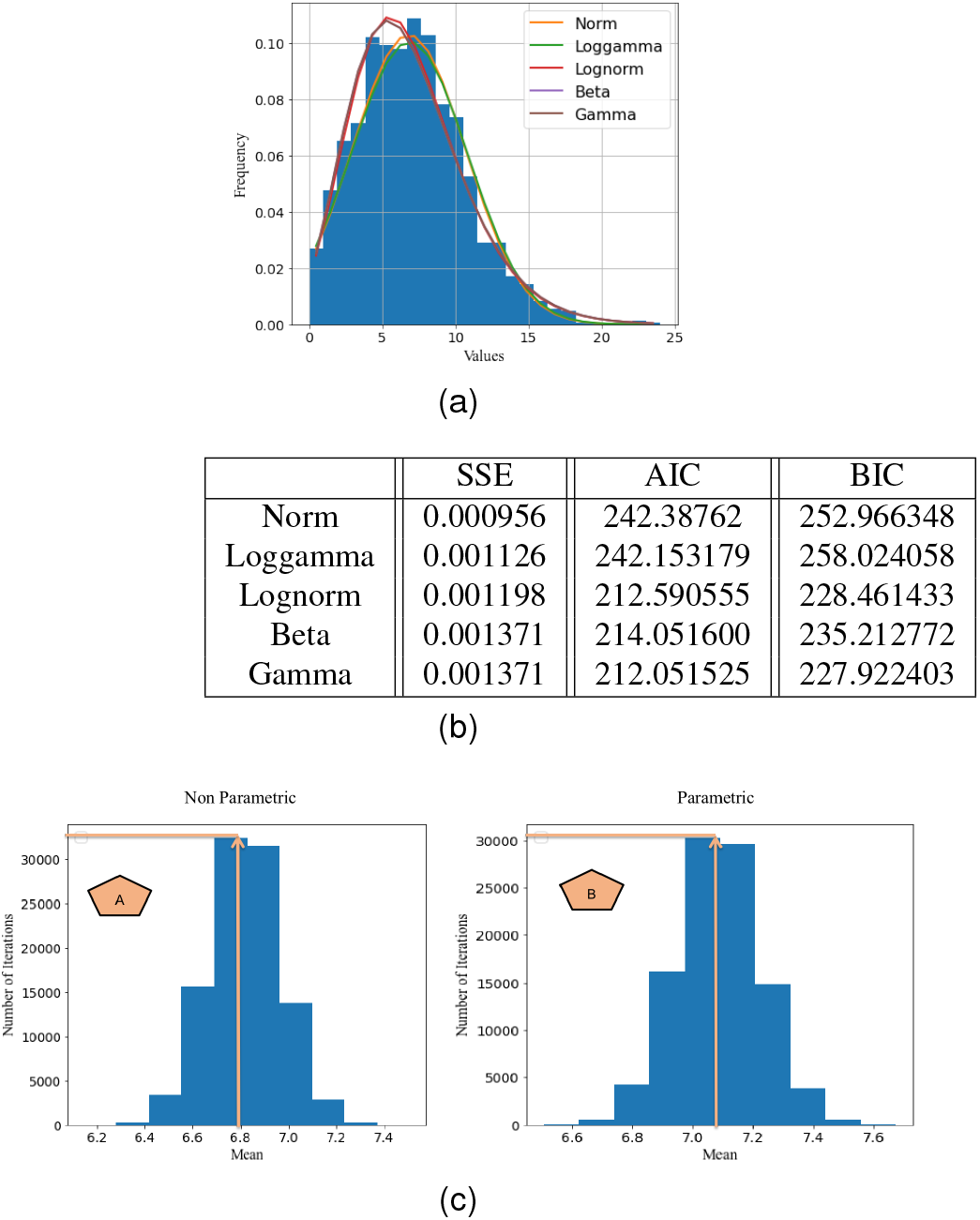
Statistical Analysis: (a) Distribution detection. (b) Ranking of distributions using SSE, AIC, and BIC. (c) Mean interval comparison between original and simulated signals. Observations: A - Observed mean matches original at max iteration; B - Observed mean exceeds original at max iteration.

### B. Statistical analysis

We randomly selected samples *B* times, each with a size of *n* from the *P* dataset or from an estimated distribution based on its parameters. The *P* dataset refers to the original population or dataset from which the samples are drawn. This could represent, for example, a collection of DNA sequences or any dataset relevant to the study [10]. For each of these *B* set, we computed key statistics such as the mean and variance. These statistics represent the characteristic distribution of samples. Next, we aggregated the statistics from all *B* samples to calculate the overall mean and overall variance. These aggregated values converge to the true mean and variance of the original *P* dataset, providing a reliable estimate of its statistical properties. To evaluate the accuracy of these statistical estimates, we used parametric and non-parametric bootstrap sampling methods [28]. Bootstrap sampling involves repeatedly resampling from the dataset (with or without replacement) to assess the variability of the estimates. We calculated the Confidence Interval (CI) for the estimates by conducting multiple iterations of this process. For constructing the CI, we preferred the standard error method over the percentile method, as the distribution of the statistics closely resembles a normal distribution, making the standard error method more appropriate. Finally, to determine the best sampling method, we performed a comprehensive evaluation using hypothesis testing, histogram analysis, distribution analysis, confidence intervals, and the range of mean values (as shown in Supplementary S2 Figure 2). This multi-faceted approach ensures that the chosen sampling method provides accurate and reliable estimates of the dataset’s statistical properties.

The statistical test was performed with data available from the experiment in [10] to test the hypothesis of whether original and sampled populations have equal means (Supplementary S2 Table 2). From Table I, it is observed that the non-parametric method has a higher p-value than the parametric method, and both are not statistically significant (P*>*0.05), indicating strong evidence for the null hypothesis.

**TABLE I.** Welch t-test result

| Stats | Parametric and original sample count | Non-Parametric and original sample count |
| --- | --- | --- |
| t-statistic | -1.56 | -0.26 |
| p-value | 0.12 | 0.79 |

Histograms created for the original and simulated samples using both parametric and non-parametric sampling models appear similar, but the fitted distributions differ. We noticed that the normal distribution, which represents the original sample data from the experiment, is the primary distribution in the non-parametric model. In the parametric samples, the log-normal distribution is detected as the primary distribution. This once again highlights the advantage of using non-parametric methods over parametric ones to approximate the copy count distribution (Supplementary S2 Figure 3).

We conducted further analysis to determine the better sampling method through Confidence Interval (CI) and mean range estimation. A good sampling method should work with a lower CI. Through trial and error tuning experiments, we identified that non-parametric bootstrap sampling works with a 90% CI and a mean range of [6.6733, 6.9696], while parametric bootstrap sampling works with a 95% CI and a mean range of [6.8158, 7.3589] (Supplementary S2 Table 3).

As shown in Fig. 3c, the original mean value of 6.8 is centered in the case of non-parametric fitting and shifted to the right in parametric fitting. This means that the same mean value as the original is generated more times in non-parametric sampling, demonstrating that non-parametric fitting performs better in terms of approximating the real data.

### C. Implementation of copy distribution for various coverages values

After approximating the copy count distribution from the real dataset at a specific coverage level, we equipped the simulator with an option to generate copy count distributions under various coverage conditions. To simulate additional sequence copies from a small sequence population, sampling with replacement is performed, where the probability of selecting a sequence is proportional to the inverse of its coefficient of variation. Specifically, we define the inverse CV for each sequence as 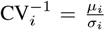, where *µ*_*i*_ and *σ*_*i*_ denote the mean and standard deviation of the observed counts, respectively. This approach reflects a reliability-aware strategy: sequences with higher abundance consistency (i.e., lower relative variability) are sampled more frequently. Such a method is particularly appropriate in simulation scenarios where coverage stability, amplification robustness, or confidence in fragment abundance are critical factors. We iterated the simulation 1000 times and retained only those sample sets whose inverse CV values fell within a target range, thereby ensuring coverage reliability.

From the filtered sets, the sample set with the lowest inverse coefficient of variation (i.e., highest reliability and consistency in sequence representation) was selected. At higher coverage values, non-parametric sampling led to a narrower range of inverse CV values ([1.760, 1.769]), indicating minimal variation across replicates and suggesting stable sampling behavior. In contrast, parametric sampling introduced a broader range ([1.76, 2]), reflecting increased variability in the representation of sequence copies. At lower coverage levels, the trend persisted—non-parametric sampling yielded relatively tighter inverse CV ranges ([1.20, 1.77]) compared to parametric sampling, which exhibited a wider distribution ([1.00, 1.77]). These observations imply that narrower inverse CV ranges correspond to more consistent sampling outcomes with reduced variability, particularly in the context of replacement-based sampling strategies (Supplementary S3 Figure 4).

### D. Preprocessing module and computation of CI and Inverse CV for all distributions

Based on the previous findings, we proposed a preprocessing algorithm as shown in Fig. 4. From the real-time hdf5 file of [10], 10,000 samples were extracted, and the repetition of each encoded input sequence in sample set A was recorded. If an encoded sequence is not mapped with any of the samples, it is termed as lost with a count of 0 in sample set A (Supplementary S4 Figure 5). The non-parametric bootstrap sampling is iterated 10,000 times from sample set A to extract specific sizes of samples. As a result, we obtained 10,000 estimated sample sets A1, A2, … A10000 with specified sample sizes consisting of different count values. First, the mean of these sets is computed, resulting in 10,000 means. Next, we averaged the resulting 10,000 means and calculated the standard deviation value. Then, using these outputs and setting the percentile at 90 using the standard error method, we computed the mean range through CI calculation for all the distributions. Next, we determined the tuned inverse CV. If the coverage value set by the user is the same as the actual mean (i.e., the mean of sample set A), the process is stopped. However, if the coverage value is greater or less than the original mean, the probability-based sampling is executed 1000 times, and the inverse CV is tuned with values as shown in Table II. Finally, we extract one sample set after filtering.

**TABLE II.** Confidence interval (ci) and Inverse coefficient of variation (cv) with respect to coverage

| Distribution | CI of mean | Inverse CV (equal coverage) | Inverse CV (higher coverage) | Inverse CV (lower coverage) |
| --- | --- | --- | --- | --- |
| Normal | [6.662, 6.982] | [1.610, 1.630] | [1.610, 1.630] | [1.100, 1.770] |
| Log-gamma | [13.354, 13.931] | [1.813, 1.815] | [1.750, 1.850] | [1.100, 1.800] |
| Beta | [11.110, 11.695] | [1.490, 1.500] | [1.490, 1.500] | [1.100, 1.500] |

**Fig. 4.**
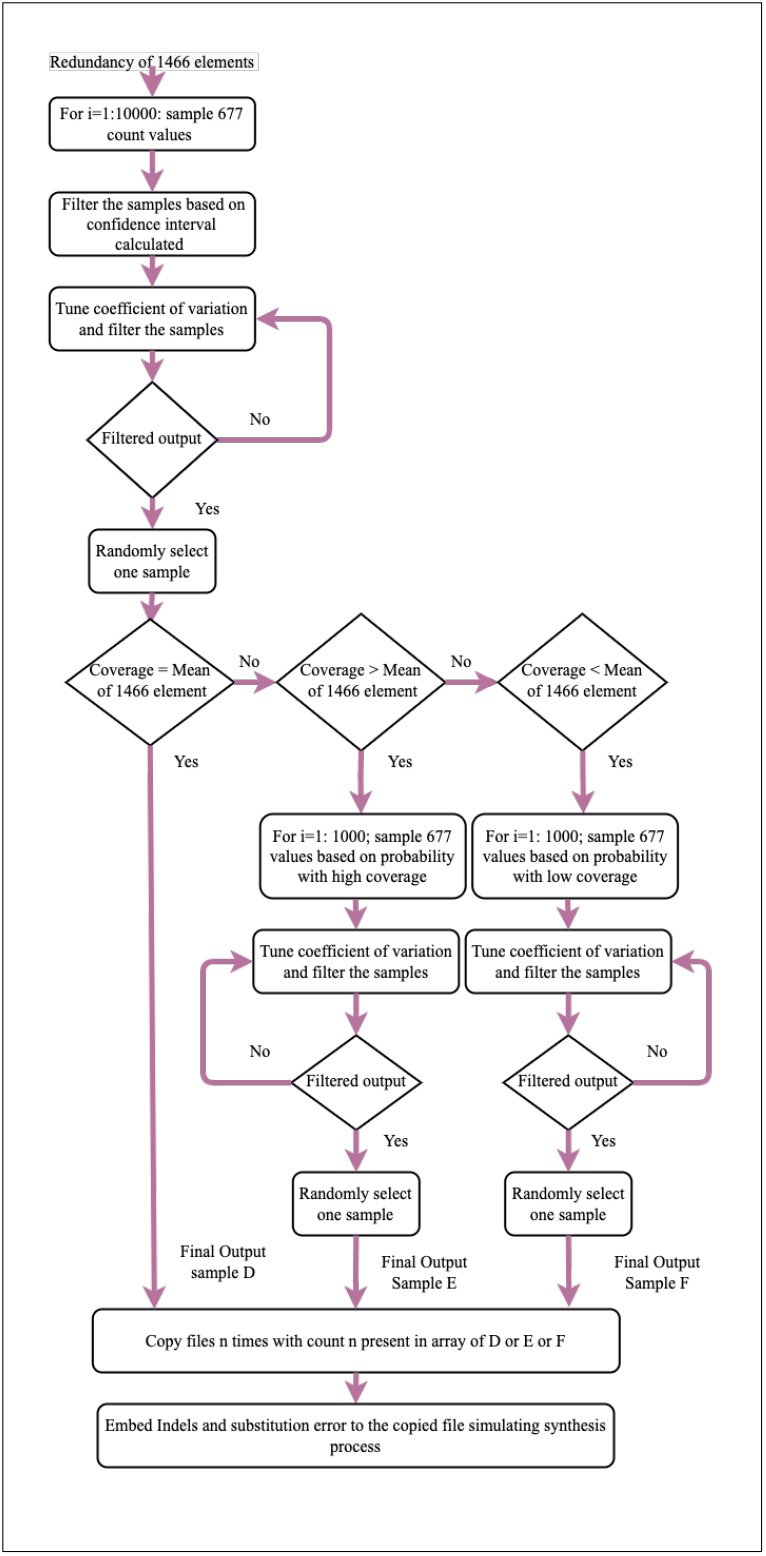
Flowchart for integrating sequencing coverage and error rate

### E. Integration of the Preprocessing module in DeepSimulator

In the sequence generator module described in DeepSimulator [26], each read from an input file (which contains ‘k’ reads) is distributed into ‘k’ separate files. These individual files are then sent to the signal generator, where a signal is generated for each read. This signal is subsequently passed to the basecaller, which converts it into a base-called sequence. DeepSimulator generates one output sequence for each input sequence, reflecting a single read. In contrast, the proposed simulator introduces a more realistic approach by generating multiple copies of each input sequence. These copies are created based on a sequencing coverage distribution, which is defined in Section III.A and can be adjusted using a tunable parameter. This approach simulates the redundancy introduced during the processes of DNA synthesis, PCR amplification, and sequencing. The generated copies are embedded with errors—insertions (I), deletions (D), and substitutions (S)—that are randomly distributed, mimicking the types of errors commonly observed in practical DNA data storage systems.

In this study, we simulated nanopore sequencing reads by integrating error profiles from DNA chemical synthesis, PCR amplification, and sequencing processes. Specifically, we applied substitution, deletion, and insertion probabilities of 0.4% (0.004), 0.85% (0.0085), and 0.05% (0.0005), respectively, to model cumulative errors arising primarily from synthesis and PCR, consistent with empirical observations [30]–[32]. These values yield an aggregate error rate of approximately 1.3%, which is substantially lower than 10.5% error rate typically reported for raw Oxford Nanopore Technologies (ONT) sequencing reads [33], [34]. To capture the higher error profile inherent to Nanopore sequencing, we incorporated DeepSimulator—a well-established nanopore read simulator that reproduces ONT’s raw read error rates—into our workflow. By integrating synthesis- and PCR-derived errors with DeepSimulator’s nanopore-specific error model, we developed the D2Sim model, which realistically reflects the layered sources of error from synthesis through to sequencing. This combined approach allows simulation of nanopore reads that encompass biologically relevant errors introduced during DNA preparation as well as the characteristic noise and inaccuracies of nanopore sequencing and basecalling, thereby providing a comprehensive and experimentally supported framework for in silico nanopore data generation.

As a result, the generated copies are embedded with errors that are randomly distributed, mimicking the types of errors commonly observed in practical DNA data storage systems. In addition, at the receiver end, each original sequence is no longer represented by just one copy but by several redundant copies. This redundancy reflects the real-world scenario where multiple copies of a sequence are produced due to the amplification and sequencing processes. The error-embedded copies provide a more accurate simulation of the challenges faced in DNA data storage. To further enhance the simulation, the proposed preprocessing module (illustrated in the dotted box of Fig. 2) allows users to select from three different distributions—normal, log-gamma, and beta—to model sequencing coverage. Additionally, the module supports customized coverage levels of up to 62 (as detailed in Supplementary S5). This flexibility ensures that the simulator can adapt to various experimental setups and accurately reflect the behavior of DNA sequences under different conditions. This approach, derives distribution using real experimental data from [10], provides a more comprehensive and realistic simulation framework compared to DeepSimulator, making it a valuable tool for DNA data storage research.

## IV. Validation of simulated signals

To validate the accuracy of D2Sim, we used a benchmark real-time experimental dataset [10]. Multiple outputs generated by D2Sim for a given encoded sequence of the experimental dataset were compared to multiple real outputs from the Nanopore sequencer of the experimental dataset. Both cross-correlation and DTW analyses confirmed that D2Sim exhibited closer temporal and morphological alignment to the real signals than the existing Deepsimulator. For each pair of signals, we evaluated the lag required to best align them. A smaller lag value indicates a closer match in temporal structure. The results showed that the proposed simulator, D2Sim, exhibited significantly lower sample difference deviations compared to DeepSimulator.

DTW distances further confirmed that D2Sim output signals were closer in shape to experimental signals, demonstrating improved simulation fidelity. The two signals were stretched onto a standard set of instants such that the sum of Euclidean distances between the corresponding points was minimized. By doing this, the visual representation of the signal became similar, as shown in Fig. 5a. Please refer to Supplementary S6 Figure 7 for correlation between real signals and simulated signals.

**Fig. 5.**
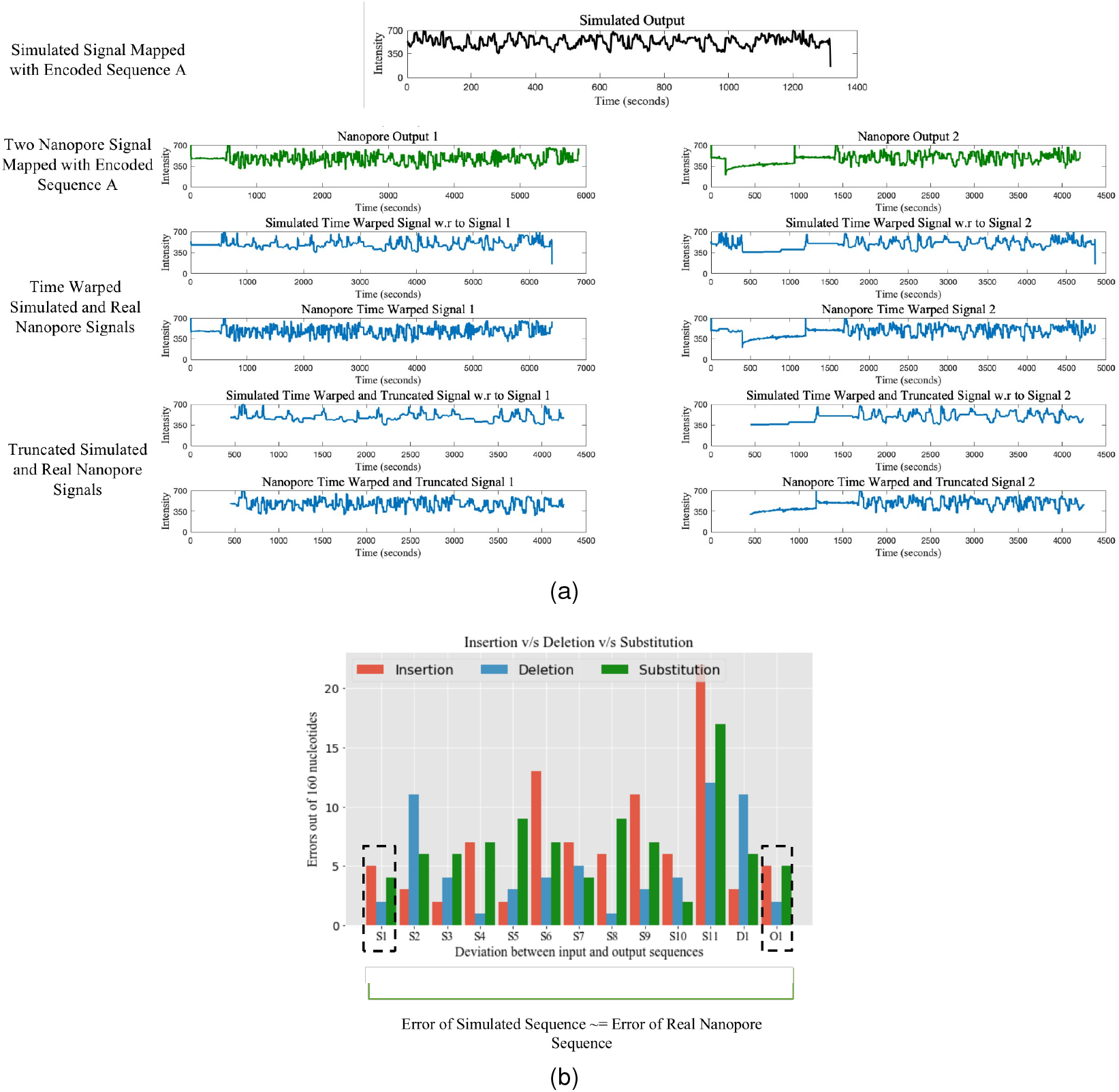
Validation: (a) Analysis of signals. Simulated signals from the proposed simulator having the least sample difference (with respect to 2 real Nanopore signals) are taken and represented by a pictorial representation after performing DTW and cropping. (b) Biomolecular Errors in Nanopore Sequencer (O1), proposed simulator D2Sim (S1-S11), and DeepSimulator (D1).

The characterization of errors and biases is performed by aligning sequencing reads/signals to respective references/encoded input sequences. In the designed simulator, the sequences are embedded with certain insertion, deletion, and substitution rates, whereby the error rates at the receiver are closer to real DNA data storage. In our study, we utilized a total of 14 sequences for validation, comprising 10 encoded input sequences detailed in the supplementary material (Supplementary S6 Table 4a) and an additional 4 original output nanopore signals in this paper.

As shown in Table III, we took four encoded input sequences and correspondingly four original output Nanopore signals from [10] for validation. Each of the input sequences was processed in DeepSimulator, producing one simulated output per sequence. Following that, the sample difference between the simulated and the original nanopore signal was calculated through signal correlation. Similarly, we have extracted 11 simulated copies per sequence using the proposed DNA storage channel simulator and computed the sample difference between the simulated and original Nanopore signal. From the table, it is found that DeepSimulator generates a signal with a higher sample difference compared to the original output signal. In contrast, the proposed simulator D2Sim produces certain output signals with lower sample differences than the original Nanopore signal. The signals generated by the proposed simulator have 16.7% to 88.7% lower sample differences than the signals obtained from DeepSimulator, enabling scientists to handle varied error scenarios efficiently. In Supplementary S6 Figure 6, the visual representation of the signal is shown for signals having sample differences of 86 and 27, as underlined in Table III.

**TABLE III.**
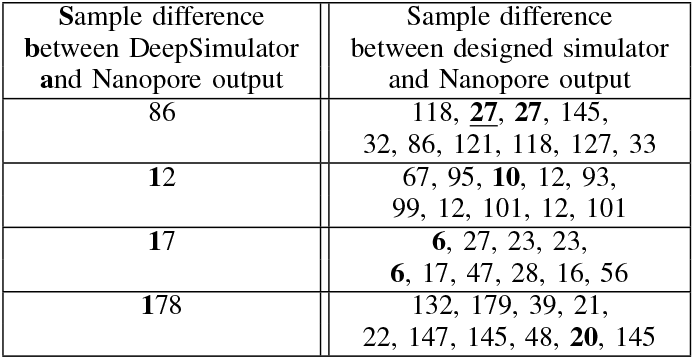
Sample difference calculation between Nanopore signal and simulated signals (having sample length of around 4000 due to biomolecular error after DTW) mapped to input encoded sequence of length 160

| Sample difference between DeepSimulator and Nanopore output | Sample difference between designed simulator and Nanopore output |
| --- | --- |
| 86 | 118, <u>27</u>, <u>27</u>, 145, 32, 86, 121, 118, 127, 33 |
| 12 | 67, 95, <u>10</u>, 12, 93, 99, 12, 101, 12, 101 |
| 17 | <u>6</u>, 27, 23, 23, <u>6</u>, 17, 47, 28, 16, 56 |
| 178 | 132, 179, 39, 21, 22, 147, 145, 48, <u>20</u>, 145 |

In addition, we also observed that a signal and its redundant copies read by the Nanopore sequencer have a certain sample difference of 286 (as shown in Table IV), which can be mapped to the same encoded input sequence. It further strengthens the validation of the proposed simulator as signal copies generated from the proposed simulator also differ significantly in sample difference. Now, using two real Nanopore output signals of the same original sequence, validation is carried out by estimating the sample difference between 11 simulated outputs and the two copies. The differences between the simulators and the Nanopore outputs were computed, where we observed that the proposed simulator can produce outputs having 40 and 5 samples difference from that of the original Nanopore signals, while for DeepSimulator it is 2681 and 5. To illustrate the similarity between the simulated signal and the two Nanopore signals, we display the second simulated copy that has the least overall sample difference from the real Nanopore signals (40 with respect to one real copy and 48 with respect to the other, as underlined in Table IV), after performing DTW and cropping in Fig. 5a. Additionally, comparison between multiple nanopore real signal and multiple real signals corresponding to same encoded sequence is calculated in Table 5 of Supplementary material.

**TABLE IV.** Sample difference calculation between two Nanopore signals and simulated signals (having sample length of around 4500 due to biomolecular error after DTW) mapped to input encoded sequence of length 160

| Difference b/w 2 Nanopore signals mapped to same encoded sequence : 286 | Sample difference between Deep Simulator and Nanopore output | Sample difference between designed simulator and Nanopore output |
| --- | --- | --- |
| Redundant Real Signal 1 | 2681 | 2680, <u>40</u>, 2694, 2681, 50, 71, <u>40</u>, 2681, 2687, 2694 |
| Redundant Real Signal 2 | 5 | 18, 48, 64, <u>5</u>, 61, 38, 47, <u>5</u>, 59, 40 |

In the supplementary material (Table 4a), to assess the fidelity of simulated signals, we computed the sample difference (post-DTW alignment) between real Nanopore signals and outputs from both D2Sim and DeepSimulator, using ten input-encoded sequences. Each real signal was compared against ten simulated copies from D2Sim and one copy of DeepSimulator. Across all cases, D2Sim consistently demonstrated lower sample differences in a majority of comparisons. In particular, in 6 out of 10 real signals, at least one D2Sim-generated signal had a lower sample difference than any DeepSimulator counterpart. The number of D2Sim simulated copies outperforming DeepSimulator ranged from 3 to 7 per sequence. In one case, a D2Sim copy exactly matched the real Nanopore signal (zero sample difference). For the remaining 3 signals, DeepSimulator achieved a lower sample difference. These findings suggest that D2Sim produces signal traces with higher temporal fidelity, better preserving the characteristics of real Nanopore signals. This supports its utility for downstream evaluation tasks requiring signal-level realism, such as base- caller benchmarking or signal-model training.

A complementary evaluation in the supplementary material (Table 5) focuses on signal-level mapping accuracy, where multiple real and simulated signals were compared for each of 10 encoded sequences. D2Sim achieved exact signal-level matches with real reads in up to 50% of simulated outputs, with exact match percentages varying across sequences ([0, 5.3, 8.3, 12.5, 13.3, 37.5, 50]). In 7 out of 10 input sequences, D2Sim produced simulated signals that were more similar to real reads than those from DeepSimulator, with improvements in top-match accuracy ranging from 12% to 83%. Conversely, DeepSimulator performed better in 3 cases, showing top-match advantages in the range of 8% to 50%. These findings collectively demonstrate that D2Sim captures signal-level distortions and variability more effectively, resulting in outputs that are statistically more similar to actual nanopore read patterns.

As reported in Table 4b (Supplementary), the designed simulator generates signal alignments with a markedly broader distribution of temporal lags (sample differences) compared to DeepSimulator. While DeepSimulator consistently outputs a fixed lag per signal, the designed simulator produces multiple signal realizations with minimum-to-maximum lags varying substantially. This high variability reflects the stochastic nature of real nanopore sequencing, where current signal durations can fluctuate due to complex physical interactions, molecule speed variation, and instrument noise. The ability of the designed simulator to produce such a diverse signal lag distribution offers a more biologically faithful alternative to deterministic approaches like DeepSimulator, making it more suitable for alignment tools, basecallers, and signal analysis methods under realistic variability conditions.

Further validation is performed on the biomolecular errors between the input encoded sequence and sequences from real Nanopore sequencer, DeepSimulator, and the proposed D2Sim as shown in Fig. 5b. We observed that the proposed simulator produces an output having comparable biomolecular errors (Insertion: 5, Deletion: 2, Substitution: 4) to the Nanopore sequenced output (Insertion: 5, Deletion: 2, Substitution: 5), whereas DeepSimulator produces highly deviated errors, especially in deletions (Insertion: 3, Deletion: 11, Substitution: 6). Also, one important observation is that the primer sites are affected the most by errors compared to the center. For reference, Supplementary S6 Figure 8. From the validation performed, it is observed that the proposed simulator D2Sim better simulates the sequences/signal as generated in a Nanopore sequencing-based DNA data storage.

## V. Discussion

The input to a DNA storage channel is a multiset of M synthesized DNA molecules, and the output is a multiset of N reads sampled from this pool. This process is inherently noisy due to synthesis, storage, and sequencing errors—primarily insertions, deletions, and substitutions. Furthermore, not all molecules are equally likely to be sampled during sequencing; the sampling distribution, which reflects redundancy, significantly impacts the observed molecule set. This distribution was empirically modeled in our work based on prior experimental data.

A limitation reported in prior simulators [24], is their inability to accurately model the standard deviation of error rates—only the mean is tunable. As a result, real nanopore sequences often show higher variability in error compared to simulated data. However, we found that this discrepancy did not significantly affect the reliability of our simulation results either. That said, it remains an area for refinement. Additionally, DeepSimulator was found to inaccurately prioritize substitution and insertion errors over deletions [25]. Moreover, it tends to place the same error types at the same positions across reads. In contrast, our simulator introduces positionally variable errors, better reflecting experimental observations.

San Antonio et al. [24] and Hamoum et al. [25] modeled the error distributions for synthesis, PCR, and sequencing individually for nanopore based DNA storage. In contrast, our approach focuses on modeling overall coverage bias and aggregate error rates after sequencing using real experimental datasets. Sequencing errors for Nanopore are already captured by DeepSimulator, and our simulator therefore encodes only the synthesis and PCR-related error rates. These are configurable and can be adapted based on use-case.

While we do not model the positional distribution of error rates (as done in [19] for Illumina based storage, and [24] for Nanopore based storage), our simulator captures the overall distributional behavior critical for evaluating read-level variability and dropout. Our primary aim is to quantify and reproduce coverage bias, especially for long reads where Nanopore provides extended read lengths (*>* 1000 bp) compared to Illumina’s few hundred base pairs. The error probabilities in our simulator are hardcoded but can be adjusted depending on the experimental setup. While our simulator primarily focuses on modeling coverage bias, sequencing errors for Nanopore are handled externally via DeepSimulator.

Although different synthesis techniques exhibit varied error distributions, several studies have reported repeatable trends across platforms as per [17], which can guide future integration of specific error models into our framework at each phase. According to [17], insertion and deletion errors predominantly arise during synthesis, while substitution errors occur across all stages. They modeled synthesis errors with a uniform distribution and PCR bias with a long-tail distribution, which was confirmed using Illumina-sequenced experimental datasets. Their analysis demonstrated that synthesis and PCR-induced molecule maldistribution often leads to dropout, i.e., sequences that are not observed at the output. In our work, we addressed these issues in the context of Nanopore sequencing, but not Illumina. Their approach of modeling the distribution at each process step can be extended to our simulator as well, thanks to its modular architecture, allowing flexible integration of different error and sampling models.

Also, previous work has quantified insertion, deletion, substitution, and coverage bias using Illumina [35], mapping these effects across the layout and along oligonucleotide positions. These studies revealed that synthesis and photochemical efficiencies are not uniform, and error rates vary spatially across synthesis arrays—a critical insight for optimizing light-directed DNA fabrication. While we currently simulate uniformly applied base error rates, the model can be expanded to include position-specific or sequence-context-dependent error profiles, which have been shown to significantly affect retrieval fidelity, particularly for shorter or incomplete oligos.

Importantly, prior studies, such as Eva Gil-San Antonio et al. [24] and Gimpel et al. [19], have noted that error rates tend to increase near the ends of synthesized oligonucleotides, especially deletions. Our simulator captures this trend, particularly in scenarios where primer binding sites are present or absent, allowing for visual comparison and validation of such positional error patterns.

Furthermore, GC content was not explicitly modeled in our simulator, although high GC content is known to increase PCR error rates and sequence dropout [17], leading to GC-dependent coverage variation. While GC content constraints were not specifically studied in our current analysis, we acknowledge that they may influence dropout rates and error variation, and this remains a direction for future work.

To address the concern regarding sample size for performance evaluation, we emphasize that our study utilized 166 real nanopore signals and 237 simulated signals, corresponding to 23 unique input encoded sequences (including validation covered in this paper and supplementary material). This dataset enabled a robust assessment of signal-level similarity, using metrics such as Dynamic Time Warping (DTW) and cross-correlation, which are critical for validating temporal and amplitude fidelity in simulated signals. In addition to signal-level validation, we performed a focused error rate analysis (insertions, deletions, and substitutions) on 10 simulated sequences. This included comparisons between our simulator and DeepSimulator. And also our simulator generated sequences with and without primers. This targeted evaluation provided insight into how different simulator settings and the presence of primers impact error characteristics, without requiring exhaustive simulation of the entire dataset.

In the supplementary material (Tables S2–S4), we provide a comparative analysis of signal-level similarity using Dynamic Time Warping (DTW) sample differences between real nanopore outputs and the signals generated by D2Sim and DeepSimulator. These analyses show that D2Sim produces signal outputs that more closely resemble real nanopore signals in a majority of cases, with between 3 to 7 out of 10 simulated signals exhibiting lower sample differences compared to DeepSimulator. In one case, an exact signal match was observed between a real and a simulated output. While a formal statistical characterization (e.g., mean and standard deviation of differences) was not the focus of the current work, our findings demonstrate that D2Sim more consistently captures real signal variations than existing simulators. Future work will extend this analysis to include statistical distributions of signal-level metrics for deeper validation.

Although many errors are introduced stochastically, preserving signal-level characteristics is essential for downstream tasks such as basecalling and alignment. We hypothesize that a reliable simulator should introduce realistic errors without distorting the temporal dynamics and statistical properties of the raw signal. This includes maintaining fidelity in signal shape, amplitude range, and local fluctuations, which are critical for learning-based tools. Although we did not explicitly validate the coverage distribution—as it was derived from empirical observations—we validated two key components:(i) Signal-level similarity, using metrics such as Dynamic Time Warping (DTW) and cross-correlation; and (ii) Error rate statistics, including insertion, deletion, and substitution rates, which were compared to experimental data.

While our coverage depth modeling is based on empirical read count distributions from real DNA storage datasets, we acknowledge that these distributions are significantly shaped by various biochemical processes, most notably the PCR amplification steps and the oligonucleotide synthesis methods. As noted by Heckel et al. [36], PCR can introduce amplification bias, resulting in the over- or under-representation of certain oligos. Furthermore, spatial positioning of oligos on array-based synthesis platforms has been shown to affect copy number variability [37], further influencing the final coverage distribution.

In our implementation, we utilize coverage data derived from sequencing results reported in [10]. While this experimental pipeline includes both synthesis and PCR amplification, specific details regarding PCR conditions or array layouts are not fully considered. As such, our modeling focuses on the observed statistical distribution of coverage (using AIC/BIC/SSE to fit candidate distributions), without explicitly modeling the underlying mechanistic steps of PCR or synthesis.

Related efforts, such as the modeling framework presented in [38], aim to quantify and mitigate distributional biases introduced by experimental steps. These studies emphasize the importance of integrating biochemical-aware models into simulation tools. Incorporating such detailed mechanistic models in future work will further improve the realism and predictive power of simulators for DNA data storage.

Recent advances in decoding algorithms and DNA storage system design have further highlighted the importance of realistic simulation tools. In particular, the work by Sabary et al. [39] introduces efficient reconstruction algorithms capable of achieving reliable decoding with as low as 16× coverage, significantly lowering the coverage threshold for practical DNA data retrieval. Similarly, Bar-Lev et al. [40] propose a deep learning-based approach that enhances the scalability and robustness of DNA storage through sophisticated coding and decoding strategies.

Although our focus in this work is not on decoding algorithms per se, the D2Sim simulator is designed to accurately model the sequencing process, capturing real-world phenomena such as coverage bias, amplification distortion, and indel errors. As such, the datasets generated by D2Sim can be instrumental for benchmarking decoding frameworks like those proposed in [39], [40], enabling realistic evaluation under practical noise and distributional constraints.

## VI. Conclusion

The design of D2Sim follows five key steps, which include error embeddings, as well as estimating the copy count distribution to accurately reflect the behavior of DNA sequences in real-world scenarios. This comprehensive approach ensures that D2Sim provides a robust and realistic simulation framework for DNA data storage research. Due to its portability, there has been high interest in using Nanopore sequencers for performing sequencing actions in DNA storage technology. We have devised a computational simulator model D2Sim to characterize the Nanopore sequencing-based DNA data storage channel. The signals generated from the proposed simulator are validated by comparing them with the real Nanopore signals through the evaluation of sample differences and biomolecular errors. An option to specify sequencing coverage size for better adaptability to different sequencing redundancy with various experimental settings is provided, which can be used to assess the efficacy of logical and sequencing/physical redundancy, thereby providing a reference for designing encoding/decoding schemes and reconstruction methods. The maximum coverage of 62x provided by the proposed simulator can be expanded in future works by using machine learning frameworks such as General Adversarial Networks.

## References Section

The supplementary document and GitHub link is *Supplementary info DNA Storage Computational Simulator* and https://github.com/subhasiny/Computational DNA Storage Channel.

## Acknowledgments

The author thanks Shubham Chandak from Standford University for discussions regarding source data and procedure followed in their experiments.

