## Supplementary file for "D2Sim: A Computational Simulator for Nanopore Sequencing based DNA Data Storage"

### S1. Detection of distribution

The real-time signal data obtained from the Nanopore sequencer after custom array synthesis and PCR are used to find the data distribution. Analysis is carried out using data obtained from 10 experiments to determine the associated probabilistic model. Specifically, in each experiment, between 733 to 1466 encoded sequences were synthesized. After sequencing, approximately 10,000 to 20,000 reads were obtained. To determine how many times each encoded sequence appeared, the authors of [10] aligned these sequencing reads to the original encoded sequences using Minimap2. This alignment step enabled identification of how many copies of each original sequence were recovered, as well as detection of losses (i.e., sequences for which no reads were recovered). We used these alignment results to quantify coverage depth per input sequence. Then, we modeled this empirical coverage distribution and evaluated its fit against several candidate statistical distributions. Model fitting was assessed using Akaike Information Criterion (AIC), Bayesian Information Criterion (BIC), and sum of squared errors (SSE). In most cases, the distributions observed in the 10 experiments are normal, beta, and log-gamma, as shown in Table 1. Among the tested distributions, the normal distribution yielded the best overall fit to the observed data. These steps are described in supplementary material.

SSE is defined as the sum of the squared differences between observations and their group's mean. Error prediction and the relative quality of each distribution can be evaluated using a factor called AIC, and the best model should have a lower AIC. Another criterion for selecting the best model is BIC, where the best model should have a low BIC. Specifically, when statistically fitting the model to the data, adding more parameters may increase the likelihood, but it may also result in over fitting. AIC and BIC measures resolve this by introducing penalty terms for the number of parameters in the model. The penalty is higher in BIC than in AIC. When the number of observations is large, AIC and BIC may produce different model fitting results. However, since BIC applies a larger penalty for complex models, it tends to result in a more accurate fit than AIC. Therefore, in our detection algorithm, based on the priority of BIC, the second priority of SSE, and finally, AIC, the best model is determined. After modeling these distributions and evaluating the best probabilistic model, two bootstrap sampling methodologies, namely parametric and non-parametric sampling, are compared, and the better method is determined.

| Exp | Code memory (m) | Code memory | Outer code Redundancy | Encoded data(A) | Sub sample after sequencing (B) | A compared with B (C) | A-C (oligo loss) | Mean | Std | Detected Distribution | Sum square error | AIC | BIC |
|---|---|---|---|---|---|---|---|---|---|---|---|---|---|
| 0 | 8 | 1/2 | 0.3 | 1466 | 20000 | 1383 | 83 | 13.642 | 9.244 | Skew norm | 0.0026 | 729.06 | 744.94 |
| | | | | | | | | | | Log gamma | 0.0028 | 915.26 | 931.13 |
| | | | | | | | | | | Norm | 0.0028 | 928.83 | 939.41 |
| 1 | 11 | 1/2 | 0.3 | 1466 | 10000 | 1442 | 24 | 6.821 | 3.871 | Norm | 0.0009 | 242.38 | 252.96 |
| | | | | | | | | | | Log gamma | 0.0011 | 242.15 | 258.02 |
| | | | | | | | | | | Log norm | 0.0012 | 212.59 | 228.46 |
| 2 | 14 | 1/2 | 0.3 | 1466 | 10000 | 1419 | 47 | 6.846 | 4.383 | Log norm | 0.0018 | 506.73 | 522.60 |
| | | | | | | | | | | Norm | 0.0019 | 854.64 | 865.22 |
| | | | | | | | | | | Gamma | 0.0022 | 521.54 | 537.41 |
| 3 | 8 | 3/4 | 0.3 | 815 | 20000 | 810 | 5 | 24.540 | 14.846 | Log gamma | 0.0017 | 1520.66 | 1534.77 |
| | | | | | | | | | | Norm | 0.0017 | 1517.10 | 1526.51 |
| | | | | | | | | | | Beta | 0.0020 | 1275.02 | 1293.82 |
| 4 | 11 | 3/4 | 0.3 | 815 | 10000 | 807 | 8 | 12.270 | 8.434 | Skew norm | 0.0013 | 735.49 | 749.60 |
| | | | | | | | | | | Beta | 0.0018 | 719.12 | 737.93 |
| | | | | | | | | | | Gamma | 0.0019 | 679.24 | 693.35 |
| 6 | 8 | 5/6 | 0.3 | 733 | 20000 | 733 | 0 | 27.285 | 20.894 | Skew norm | 0.0014 | 547.96 | 561.96 |
| | | | | | | | | | | Gamma | 0.0017 | 527.48 | 541.59 |
| | | | | | | | | | | Beta | 0.0017 | 553.86 | 572.67 |
| 7 | 11 | 5/6 | 0.3 | 733 | 10000 | 731 | 2 | 13.642 | 5.736 | Log norm | 0.0017 | 1972.53 | 1986.33 |
| | | | | | | | | | | Gamma | 0.0021 | 2069.71 | 2083.50 |
| | | | | | | | | | | Beta | 0.0021 | 2083.89 | 2102.28 |
| 8 | 14 | 5/6 | 0.3 | 733 | 10000 | 724 | 9 | 13.642 | 7.851 | Skew norm | 0.0007 | 384.94 | 398.73 |
| | | | | | | | | | | Log norm | 0.0007 | 382.48 | 396.27 |
| | | | | | | | | | | Gamma | 0.0007 | 383.61 | 397.41 |
| 9 | 11 | 3/4 | 0.2 | 752 | 10000 | 744 | 8 | 13.297 | 8.018 | Norm | 0.0020 | 615.59 | 642.78 |
| | | | | | | | | | | Log gamma | 0.0020 | 613.89 | 627.68 |
| | | | | | | | | | | Log norm | 0.0026 | 531.36 | 545.16 |
| 10 | 11 | 3/4 | 0.4 | 877 | 10000 | 864 | 13 | 11.402 | 7.572 | Beta | 0.0022 | 444.07 | 462.56 |
| | | | | | | | | | | Log gamma | 0.0025 | 485.30 | 499.16 |
| | | | | | | | | | | Norm | 0.0025 | 483.92 | 493.17 |

Table 1. Distribution of all experiments

### S2. Statistical Analysis

Bootstrap sampling (Figure 1) is used to simulate samples from a large sample set. This allows the calculation of standard errors, confidence intervals, and hypothesis testing. The population distribution can be estimated from the observed samples. Different forms of bootstrap sampling, including non-parametric and parametric methods, are evaluated using the techniques illustrated in Figure 2.

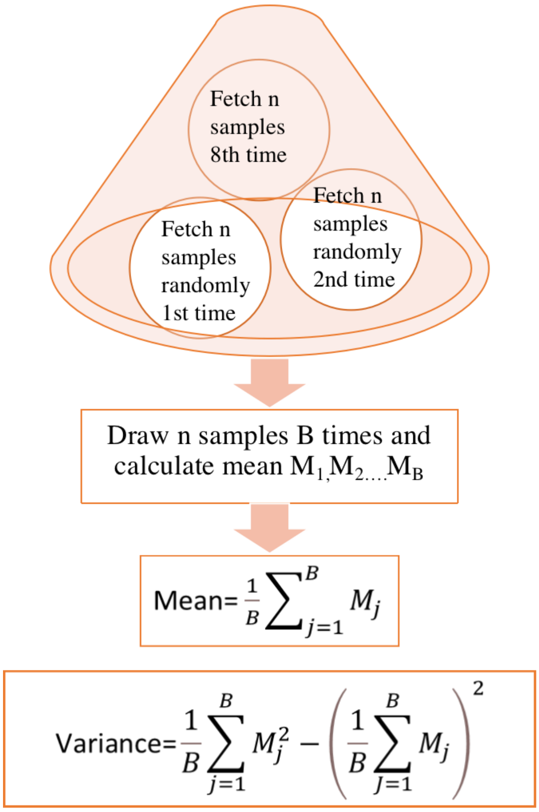


Figure 1. Bootstrap Sampling

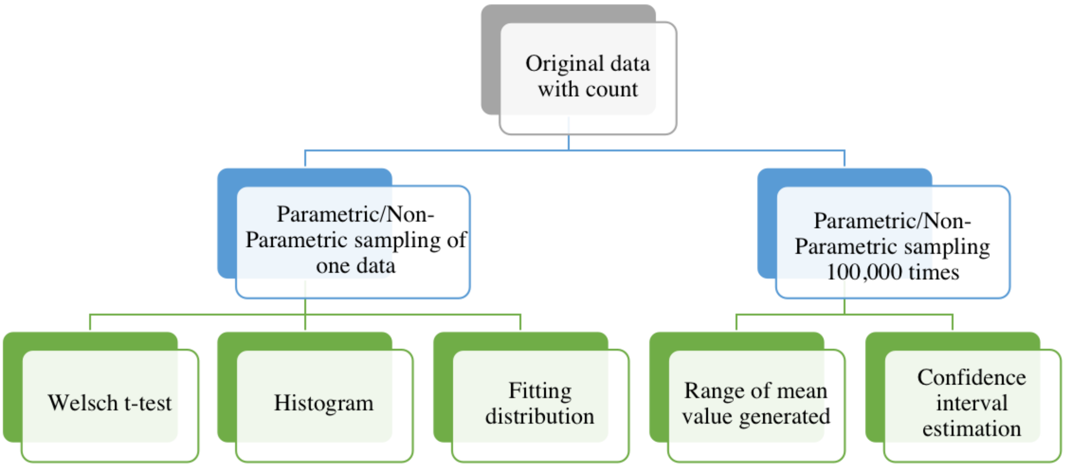


Figure 2. Statistical analysis for evaluating sampling methodology

**Welsch t-test** is an adaptation of the student's t-test for samples with unequal variance and sample size. It is performed on sampled dataset

$$t = \frac{\bar{x}_1 - \bar{x}_2}{\sqrt{\frac{s_1^2}{N_1} + \frac{s_2^2}{N_2}}} \qquad v \approx \frac{\left(\frac{s_1^2}{N_1} + \frac{s_2^2}{N_2}\right)^2}{\frac{s_1^4}{N_1^2 v_1} + \frac{s_2^4}{N_2^2 v_2}}$$

Where $\bar{x}_1$ and $\bar{x}_2$ represents the means of the original and sampled data, $N_1$ and $N_2$ represents the numbers of the original and sampled data, $S_1$ and $S_2$ represents the standard deviations of the original and sampled data, $v_1$ and $v_2$ represents the degrees of freedom from the original and sampled data $v_1 = N_1 - 1$ and $v_2 = N_2 - 1$

Table 2. Welsch t-test statistical result

| | Original samples | Parametric samples | Non-parametric samples |
|---|---|---|---|
| Mean | 6.821 | 7.085 | 6.867 |
| Standard deviation | 3.871 | 3.520 | 3.776 |
| Sample size | 1466 | 677 | 677 |
| t-statistic with original samples | | -1.5655117655280522 | -0.2588118466420289 |
| p-value | | 0.11768337368772319 | 0.7958200151981172 |

The **histogram** of the simulated dataset (consisting of 677 samples) and the original dataset is shown in Figure 3, along with the **distribution estimation** graph

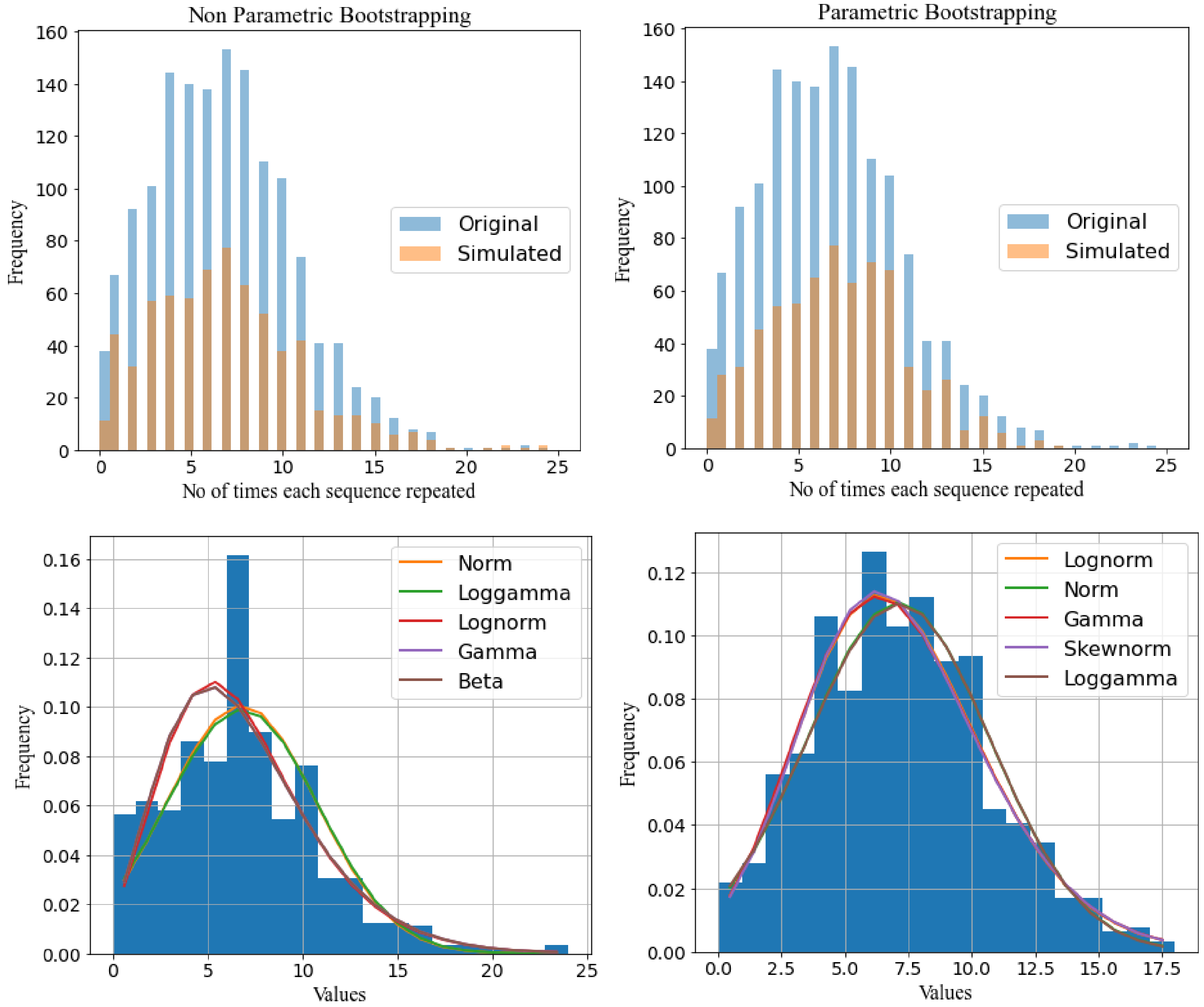


| | SSE | AIC | BIC |
|---|---|---|---|
| Norm | 0.006687 | 191.452281 | 200.487624 |
| Loggamma | 0.006793 | 192.421224 | 205.974237 |
| Lognorm | 0.007384 | 168.440369 | 181.993383 |
| Gamma | 0.007612 | 168.129164 | 181.682178 |
| Beta | 0.007612 | 170.132058 | 188.202743 |

| | SSE | AIC | BIC |
|---|---|---|---|
| Lognorm | 0.001965 | 131.296263 | 144.849277 |
| Norm | 0.002031 | 132.061052 | 141.096395 |
| Gamma | 0.002033 | 131.041024 | 144.594038 |
| Skewnorm | 0.002041 | 131.257001 | 144.810014 |
| Loggamma | 0.002063 | 134.281357 | 147.834371 |

Figure 1. Histograms of sequences and distribution

The mean range is computed by determining the best confidence interval, as shown in Table 3. If the confidence interval is narrowed further in parametric bootstrap, the next filtering step using the coefficient of variation is aborted. This means that parametric bootstrap sampling requires a greater number of bags to be sampled to determine the best sample. This decreases the probability of obtaining samples with estimated statistics like the true statistics in a single sampling process. Therefore, non-parametric sampling is preferable as it provides count values like the true statistics.

### S3. Implementation of Copy Distribution for various Coverage values

The increase in coverage value needs to be addressed as it plays a vital role in the DNA channel model when it comes to sequencing. For instance, in Experiment 1, as shown in Table 1, the coverage is 6.8x for the randomly extracted dataset (obtained by evaluating the redundant copies of experimental data). However, the proposed algorithm must be able to sequence more or fewer copies with coverage greater than or less than 6.8x. Hence, sampling with replacement is carried out based on the probability of the occurrence of each sequence when coverage varies from 6.8x. Simulated copies and the original distribution are depicted in the histogram (Figure 4).

Table 3. Confidence interval tuning using standard error method

| | Non-parametric | Parametric |
|---|---|---|
| Confidence Interval | 90% | 95% |
| Mean range formula calculated from confidence interval | [Mean - std, Mean + std] | [Mean-(1.96*std), Mean+(1.96*std)] |
| Mean range<br>Original mean: 6.821 | [6.673278, 6.969596] | [6.815854, 7.358944] |
| Coefficient of variation range,<br>Original: 1.762123 | [1.762100,1.762199] | [1.762100, 1.762199] |

After inferring results from various analyses, non-parametric bootstrap sampling is considered the best option due to various reasons, including the p-value, drift of the mean range, higher changes in the coefficient of variation compared to the original data, and a lower confidence interval.

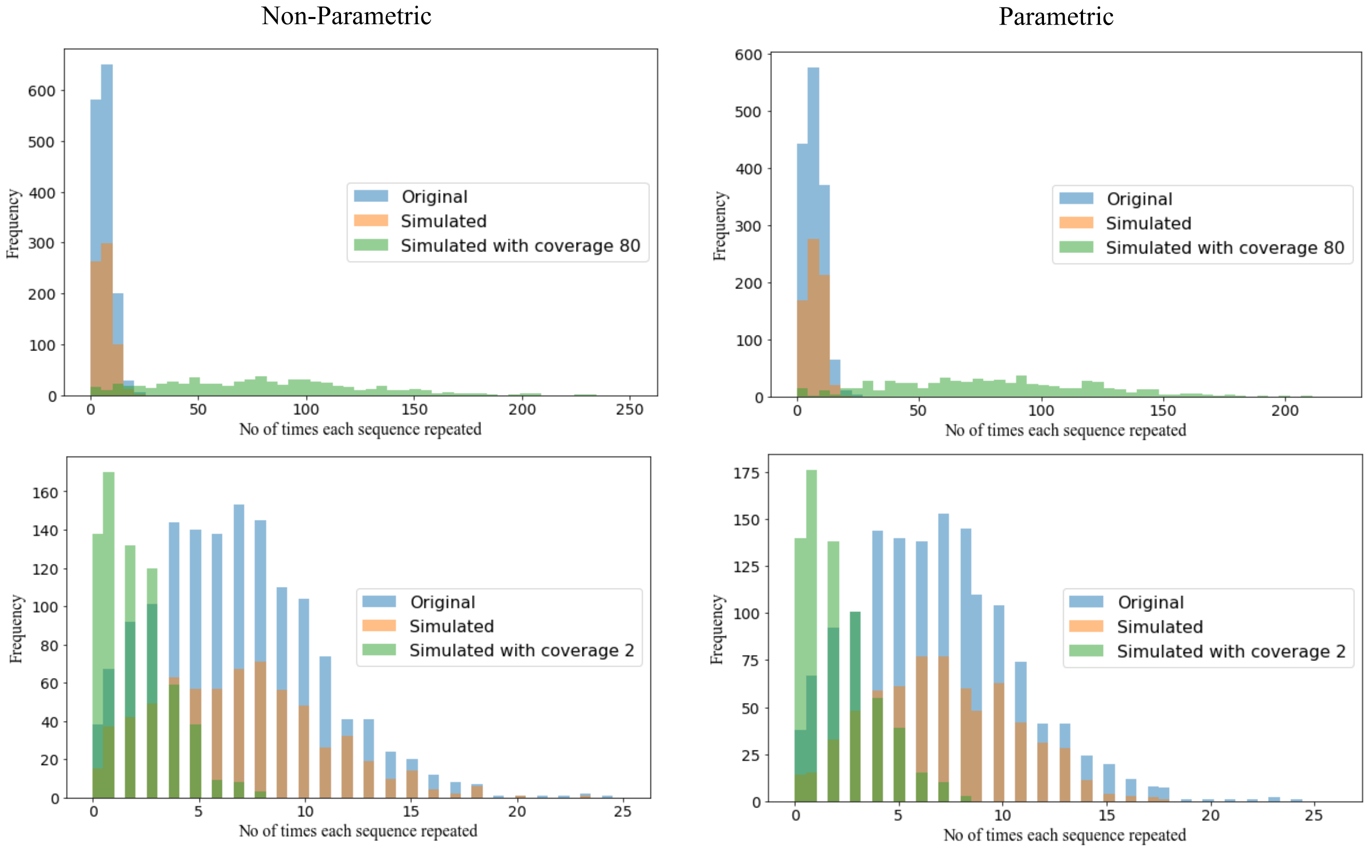

Figure 4. Histograms for different coverage

### S4. Preprocessing module and Computing Confidence Interval and Inverse Coefficient of Variations for all distributions

The algorithm for calculating the predetermined confidence interval for each distribution case (normal, beta and log-gamma) is shown in Figure 5.

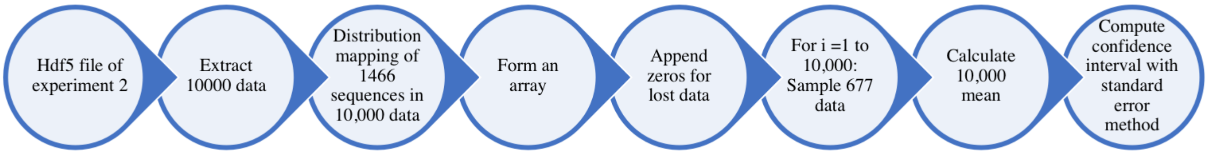

Figure 5. Computing Confidence Interval

### S5. Integration of Preprocessing module in DeepSimulator

The external parametric input is added to specify the coverage requirement and distribution

- ✧ The parameter 'Y' is added to specify the coverage (varying from 1 to 62x) in the 'dna_storage_channel_simulator.sh' file
- ✧ The parameter 'd' is added to specify the distribution (1-normal, 2-beta, 3-loggamma)

- ✧ The parameter 'n' with the value '-1' is used to avoid the splitting of reads
- ✧ Parameters 'e', 'f', 'g', and 'o' are used to specify noise, frequency, the text signal file, and the output aligned file, respectively

**Example**: / dna_storage_channel_simulator.sh –i input_677reads.txt –n –1 –d 1 –Y 60

### S6. Validation of the Proposed Simulator

We used publicly available real Nanopore sequencing data as the experimental reference, which provides encoded sequences along with their corresponding raw current signals. In parallel, we simulated signals for the same encoded sequences using both DeepSimulator and our proposed tool, D2Sim. Thus, for each ground truth sequence, we obtained (i) experimental Nanopore signals, (ii) DeepSimulator-generated signals, and (iii) D2Sim-generated signals

To validate the results of the proposed D2Sim, three analyses have been performed. Firstly, one Nanopore signal is compared with simulated signal using Dynamic Time Warping (DTW). Upon doing so, the visual representation of the signal shows maximum similarity, as depicted in Figure 6. This applies to both the simulated signals obtained from DeepSimulator and the proposed simulator. Through the synthesizing action added in DeepSimulator, the data is copied by introducing a specific set of insertions, deletions, and substitutions, making the output closer to the original Nanopore signal (where multiple reads of the same signal are obtained from its amplified pool), resulting in a sample difference of only 27 out of 4000. However, the sample difference calculated over a length of 4000 (time) between the DeepSimulator signal and the real Nanopore signal is 86, which is comparatively high.

To illustrate how DTW quantifies signal similarity between real and simulated nanopore signals, we provide a concrete example using two synthetic, sinusoidal-like sequences. This example demonstrates the alignment process and highlights the strength of DTW in handling small non-linear temporal variations between signals.

Let the real signal be:

$$X= [\sin (0.0), \sin (0.5), \sin (1.0), \sin (1.5), \sin (2.0), \sin (2.5), \sin (3.0)]$$

and the simulated signal be:

$$Y= [\sin (0.0), \sin (0.6), \sin (1.2), \sin (1.8), \sin (2.4), \sin (3.0)]$$

We computed the DTW alignment between X and Y using a standard implementation. The cost matrix is constructed by computing the squared Euclidean distance between each pair of signal points from X and Y, and the minimum-cost path through this matrix is identified using dynamic programming.

The DTW distance obtained quantifies the overall difference between the two signals after optimal alignment. The figure below visualizes the warping path that connects corresponding elements of the two signals, showing how DTW adapts to local speedups or delays in signal progression.

This example highlights the flexibility of DTW in aligning signals with local distortions and forms the basis for comparing real nanopore signals to their simulated counterparts in our evaluation framework.

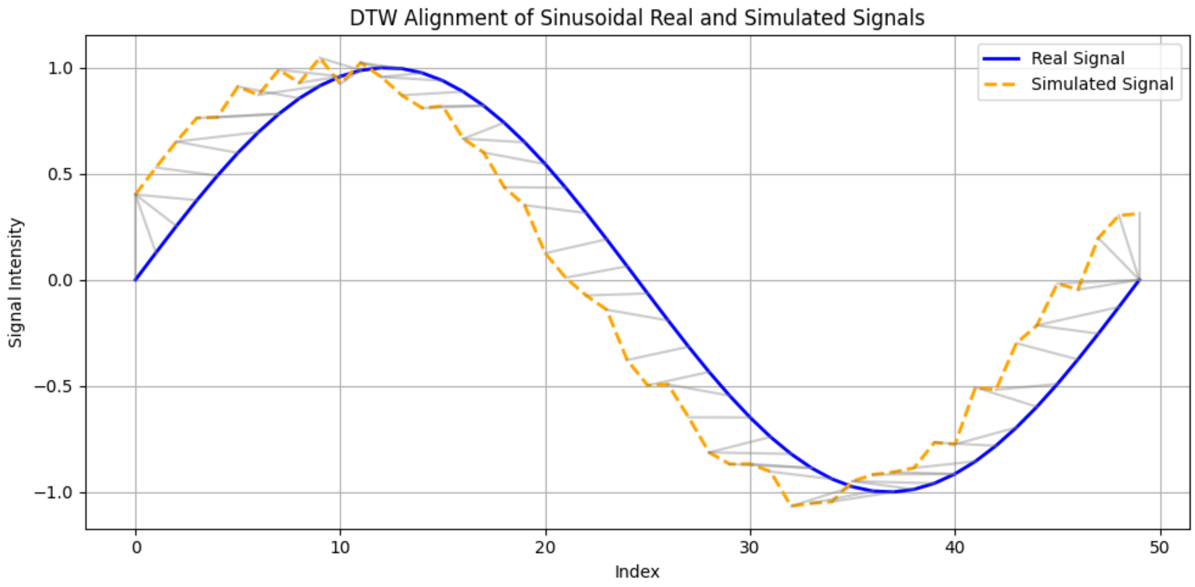

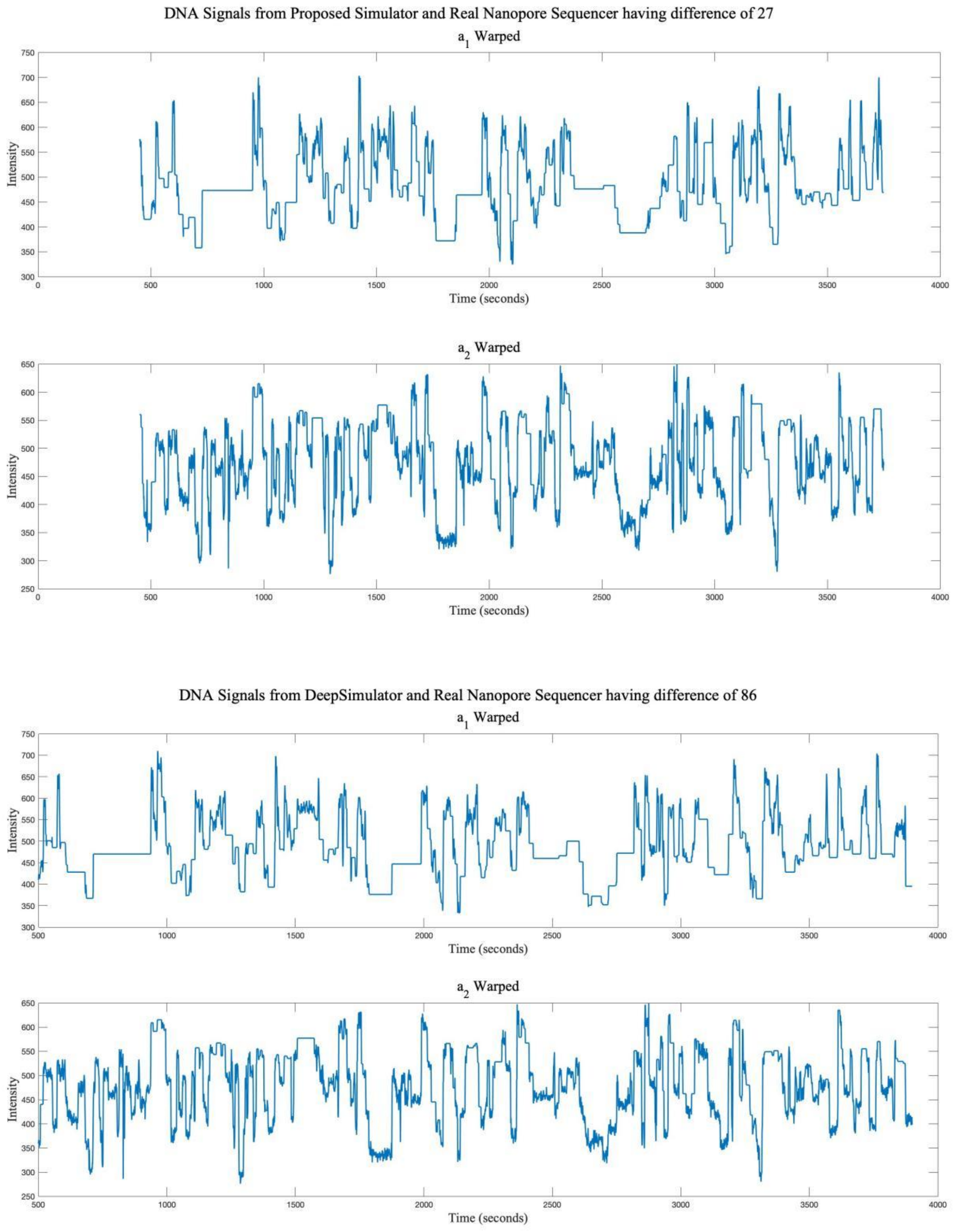


Figure 6. DWT warped signal ($a_1$ warped signal – Simulated signal and $a_2$ warped signal – Real Nanopore signal)

Secondly, two fast5 signals obtained from the original Nanopore are taken and compared with the simulated signal after dynamic time warping. Upon doing so, the visual representation of the signal appears similar. In this analysis, we are comparing the simulated output with two sets of real data and finding the difference between the two original Nanopore signals.

We computed the correlation between the signals, as shown in Figure. 5a of the main paper. The cross-correlation graph in Figure 7 reveals the maximum correlation observed between the simulated and real Nanopore signals.

Cross-correlation between Simulated signal and Real Nanopore signal 1

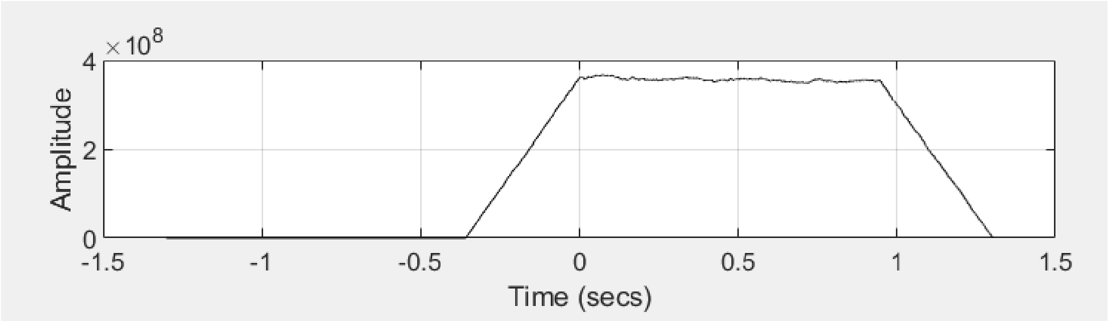


Cross-correlation between Simulated signal and Real Nanopore signal 2

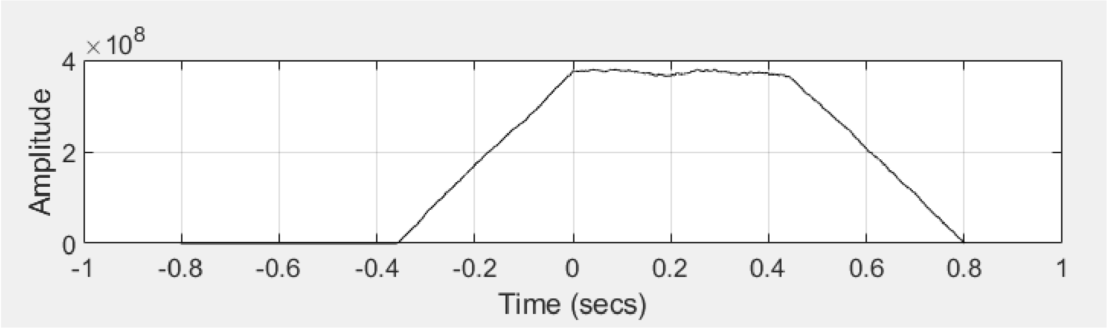


Figure 7. Cross Correlation of Signals

From Table 4, it can be inferred that the proposed simulator D2Sim simulates the signal closer to real Nanopore signal 2, with sample differences of 83, 12, 65, 5, and 89 for five signals. For the remaining simulated outputs, the sample difference is higher than 100. When comparing 10 real signals with 10 simulated copies (derived from the same input encoded sequence), one simulated copy produces output that exactly matches a real signal (signal 9). Additionally, six real signal outputs match with simulated copies having a smaller sample difference than DeepSimulator. The number of simulated copies with a smaller sample difference than DeepSimulator ranges between 3 and 7 out of 10 simulated copies. The remaining three real signal outputs match with simulated copies having a higher sample difference than DeepSimulator.

Table 4a: Observation - Sample difference calculation of two nanopore signals of the same original sequence and simulated signals (having sample length of around 4000 due to bio molecular error after DTW) mapped to input encoded sequence of length 160)

| Redundant Nanopore Output Signals | Sample Difference between DeepSimulator and Nanopore Output | Sample Difference between Designed Simulator and Nanopore Output | No. of Lowest Sample Difference in Designed Simulator than DeepSimulator |
|---|---|---|---|
| Signal 1 | 4093 | 4175,4097,4110,4096,4172,4095,4093,4093,4097 | 0 |
| Signal 2 | 1671 | 1635,1685,1681,1600,1635,1600,1671,1671,1674,1555 | 7 |
| Signal 3 | 83 | 90,34,96,96,33,96,83,83,77,90 | 5 |
| Signal 4 | 12 | 268,268,265,260,268,260,12,12,259,261 | 0 |
| Signal 5 | 147 | 288,307,319,141,288,319,147,147,302,303 | 3 |
| Signal 6 | 65 | 1793,1817,145,145,1792,145,65,65,18,140 | 3 |
| Signal 7 | 107 | 42,128,26,26,43,27,107,107,26,130 | 8 |
| Signal 8 | 5 | 29,12,5,4,29,5,5,5,9,12 | 5 |
| Signal 9 | 89 | 82,0,30,30,89,30,89,89,24,0 | - |
| Signal 10 | 251 | 293,306,280,280,285,280,251,251,280,306 | 0 |

Table 4b: Statistics Derived from Table 4a and Table III of Main Manuscript

| Signal | DeepSimulator Lag | Min Lag (Designed Simulator) | Max Lag (Designed Simulator) | Range | Std Dev | IQR |
|---|---|---|---|---|---|---|
| 1 | 4093 | 4093 | 4175 | 82 | 35.33 | 29.75 |
| 2 | 1671 | 1555 | 1685 | 130 | 43.78 | 64.50 |
| 3 | 83 | 33 | 96 | 63 | 24.20 | 16.00 |
| 4 | 12 | 12 | 268 | 256 | 106.15 | 8.00 |
| 5 | 147 | 141 | 319 | 178 | 77.39 | 123.75 |
| 6 | 65 | 18 | 1817 | 1799 | 821.08 | 1296.50 |
| 7 | 107 | 26 | 130 | 104 | 45.62 | 80.75 |
| 8 | 5 | 4 | 29 | 25 | 9.69 | 7.00 |
| 9 | 89 | 0 | 89 | 89 | 37.01 | 61.75 |
| 10 | 251 | 251 | 1280 | 1029 | 316.37 | 22.75 |
| 11 | 86 | 27 | 145 | 118 | 48.36 | 88 |
| 12 | 12 | 10 | 101 | 91 | 43.02 | 86 |
| 13 | 17 | 6 | 56 | 50 | 16.10 | 11.5 |
| 14 | 178 | 20 | 179 | 159 | 64.66 | 118.75 |

**Wide Range of Lags:** The designed simulator produces a broader range of lags compared to the DeepSimulator. This supports the claim that it can provide varied high and low lag values. **Lowest Lag Values:** The designed simulator produces lower lag values than DeepSimulator in multiple cases (e.g., Signals 2, 3, 5, 6, 7, 8, 9, 11, 12, 13, 14). **Highest Lag Values:** The designed simulator also produces significantly high lag values (e.g., Signal 10 and 12 has a max lag of 1280, much higher than DeepSimulator's 251). **High Variability:** Standard deviation (Std Dev) and Interquartile Range (IQR) are high for some signals (e.g., Signal 6 and Signal 10), showing that the designed simulator generates diverse output variations.

In Conclusion, the designed simulator (D2Sim) produces signal outputs with a broader range of temporal lags (DTW sample differences) when compared to real nanopore signals. For instance, the range of lags for a single signal span from zero to as high as 1799 samples (e.g., Signal 6), with interquartile ranges (IQR) and standard deviations significantly larger than those observed from DeepSimulator outputs. In contrast, DeepSimulator consistently produces single-valued or narrowly clustered lags (e.g., Signal 1: 4093), suggesting less signal variability.

This wider distribution in D2Sim reflects greater stochasticity and flexibility, which is expected in real nanopore sequencing due to variable molecular interactions and noise. By generating multiple outputs per sequence with differing sample alignments, D2Sim mimics the non-deterministic nature of real signal behavior more effectively than deterministic simulators. Such variability is crucial when benchmarking basecallers or signal alignment algorithms under realistic conditions.

From Table 5, it can be inferred that the designed simulator D2Sim can provide output like real sequence reads with a percentage reading from 0 to 50% [0, 5, 8, 12, 13, 37, 50]. The designed simulator can produce results more like real signals than DeepSimulator by [12, 50, 54, 59, 66, 78, 83] %. DeepSimulator produces sequences [0, 8, 15, 32, 33, 50] % which are more like real signals than the designed simulator.

Table 5: Validation of 10 sequences with r number of real signals and s number of simulated signals

| Experiment number (sequence) (r real signal, s simulated) | Exactly matched number of Designed Simulator signal to real signal (% value) | Highest match of Real signal with Designed Simulator signal compared to Deepsimulator (% value) | Highest match of Real signal with DeepSimulator signal compared to Designed Simulator signal (% value) |
|---|---|---|---|
| Exp 0 (303) [19 r, 10 s] | 1 (5.3) | 15 (78.9) | 3 (15.8) |
| Exp 6 (125) [37 r, 8 s] | 3 (8.1) | 22 (59.4) | 12 (32.4) |
| Exp 0 (1326) [12 r, 10 s] | 1 (8.3) | 10 (83.3) | 1 (8.33) |
| Exp 0 (38) [8 r, 9 s] | 3 (37.5) | 1 (12.5) | 4 (50) |
| Exp 0 (1232) [9 r, 10 s] | 0 (0) | 6 (66.7) | 3 (33) |
| Exp 8 (234) [4 r, 10 s] | 2 (50) | 2 (50) | 0 (0) |
| Exp 6 (423) [37 r, 10 s] | 5 (13.5) | 20 (54) | 12 (32.4) |
| Exp 6 (647) [24 r, 10 s] | 3 (12.5) | 13 (54) | 8 (33.3) |

Final analysis is performed by comparing simulated sequences with real Nanopore sequences in terms of bio molecular errors. With the primer, the simulated and original Nanopore outputs have a higher difference in bio molecular errors, such as insertions ranging between 12 and 18, deletions between 2 and 5, and substitutions between 3 and 11. Without the primer, the simulated and original Nanopore outputs have a lower difference in bio molecular errors, with insertions ranging between 1 and 9, deletions between 1 and 4, and substitutions between 2 and 7.

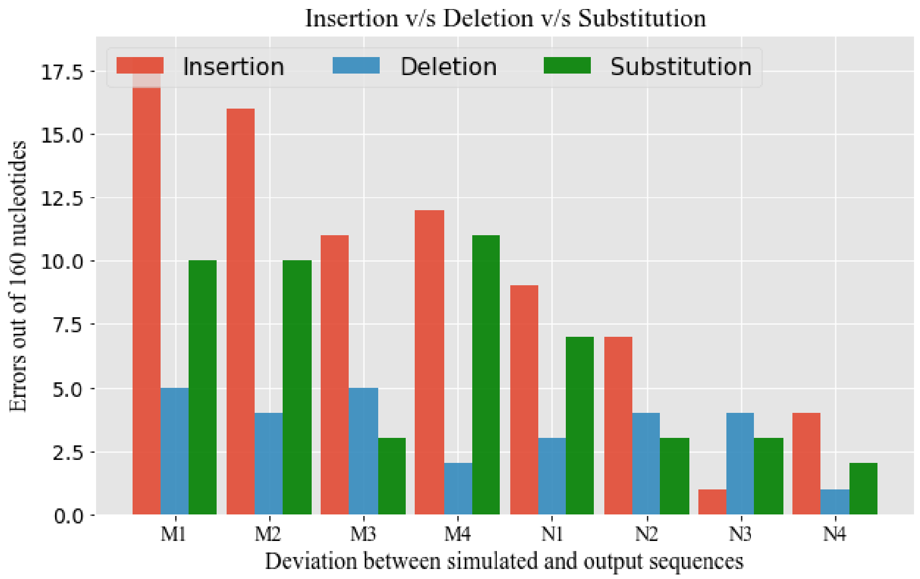


Figure 8. Bio molecular Errors with primer (M1-M4), without primer (N1-N4)

This is one of the major observations related to high occurrence of errors in the primer binding site.
